# The First Geographic Identification by Country of Sustainable Mutations of SARS-COV2 Sequence Samples: Worldwide Natural Selection Trends

**DOI:** 10.1101/2022.07.18.500565

**Authors:** Mohammadamin Mahmanzar, Seyed Taleb Houseini, Karim Rahimian, Arsham Mikaeili Namini, Amir Gholamzad, Samaneh Tokhanbigli, Mahsa Mollapour Sisakht, Amin Farhadi, Donna Lee Kuehu, Youping Deng

## Abstract

The high mutation rates of RNA viruses, coupled with short generation times and large population sizes, allow viruses to evolve rapidly and adapt to the host environment. The rapidity of viral mutation also causes problems in developing successful vaccines and antiviral drugs. With the spread of SARS-CoV-2 worldwide, thousands of mutations have been identified, some of which have relatively high incidences, but their potential impacts on virus characteristics remain unknown. The present study analyzed mutation patterns, SARS-CoV-2 AASs retrieved from the GISAID database containing 10,500,000 samples. Python 3.8.0 programming language was utilized to pre-process FASTA data, align to the reference sequence, and analyze the sequences. Upon completion, all mutations discovered were categorized based on geographical regions and dates. The most stable mutations were found in nsp1(8% S135R), nsp12(99.3% P323L), nsp16 (1.2% R216C), envelope (30.6% T9I), spike (97.6% D614G), and Orf8 (3.5% S24L), and were identified in the United States on April 3, 2020, and England, Gibraltar, and, New Zealand, on January 1, 2020, respectively. The study of mutations is the key to improving understanding of the function of the SARS-CoV-2, and recent information on mutations helps provide strategic planning for the prevention and treatment of this disease. Viral mutation studies could improve the development of vaccines, antiviral drugs, and diagnostic assays designed with high accuracy, specifically useful during pandemics. This knowledge helps to be one step ahead of new emergence variants.

**IMPORTANCE:** More than two years into the global COVID-19 pandemic, the focus of attention is shifted to the emergence and spread of the SARS-CoV-2 variants that cause the evolutionary trend.

Here, we analyzed and compared about 10.5 million sequences of SARS-CoV-2 to extract the stable mutations, frequencies and the substitute amino acid that changed with the wild-type one in the evolutionary trend.

Also, developing and designing accurate vaccines could prepare long-term immunization against different local variants. In addition, according to the false negative results of the COVID-19 PCR test report in the diagnosis of new strains, investigating local mutation patterns could help to design local primer and vaccine.

## INTRODUCTION

Genetic variations, followed by changes in phenotype level in the Sars-Cov2 virus, lead to different virus behaviors. Sars-Cov2 is a single-stranded positive-sense RNA that is approximately 30kb in length, including nonstructural (NSP) and structural proteins. The structural viral substance, including the spike (S), envelope (E), and membrane (M), as well as the nucleocapsid (N) protein that bundles the viral genome, are translated from subgenomic RNAs(1). According to research, the SARS-CoV-2 lineage, the causal agent of the pandemic COVID-19, generates one to two mutations every month as it passes through consecutive hosts, increasing infection, pathogenicity, or immune evasion(2, 3). There are six main readout frames in the virus genome open reading frame (ORF). The ORF1ab and ORF1a genes are located at the 5′ termini of the SARS-CoV-2 genome. ORF1ab contains 16 NSPs. The 3′ termini of the genome are composed of four structural proteins: S glycoprotein, E protein, M glycoprotein, and N phosphoprotein(4). The angiotensin-converting enzyme 2 (ACE2) is the primary entry receptor for S protein in SARS-CoV-2, facilitating the virus entry(5, 6). Several S mutations occurred in the appearance of B.1.1.7, B.1.351, P.1, and B.1.617.2 variants. In most cases, these mutations do not affect pathogenicity and could only affect the dissemination(5-7). N501Y, E484K, and D614G are examples of common mutations between variants(5-7). SARS-CoV-2 has over 10,000 distinct mutations compared to the reference genome(8). From the 12,509 viral genomes of SARS-CoV-2 sequences analyzed so far in the pandemic COVID-19 outbreak, the CoV-GLUE project (http://cov-glue.cvr.gla.ac.uk/#/home) has identified 5,033 amino acid replacements, with 1,687 mutations found in the S glycoprotein, 334 in the N phosphoprotein, 95 in the M glycoprotein, and 45 in the E protein(4). The D614G (S glycoprotein) mutation was thought to have originated in January 2020 in Eastern China and spread throughout the world, with the most common amino acid (AA) changes occurring in 6,855 sequences(9). So far, stable mutants have been identified for Alpha (B.1.1.7 and Q lineages), Beta (B.1.351 and descendent lineages), Gamma (P.1 and descendent lineages), Epsilon (B.1.427 and B.1.429), Eta (B.1.525), Iota (B.1.526), Kappa (B.1.617.1, 1.617.3), Mu (B.1.621, B.1.621.1), and Zeta (B.1.1.529 and BA lineages)(https://www.cdc.gov/coronavirus/2019-ncov/variants/variant-classifications.html)(10, 11). Geographically distinct SARS-CoV-2 etiological effects can be attributed to the genetic heterogeneity of SARS-CoV-2 specimens worldwide(12). The presence of so many different S protein mutations suggests numerous SARS-CoV-2 subtypes, each with its infectivity characteristics(8). In some cases, the Delta variant reduces the efficacy of therapies more than the other variants(13). Furthermore, viral mutation studies can help develop new vaccines, antiviral drugs, and diagnostic assays.

This study aimed to investigate and bioinformatically analyze stable mutations between January 2020 to May 2022 from different parts of the virus and in countries reported for the first time. By examining and analyzing with bioinformatic approaches, it has been possible to identify mutations in the coronavirus that remained stable over time and further identify each AA conversion. This study reports the top 10 mutations of each protein and the earliest reporting from each country where samples with the stable mutations occurred.

## RESULT

According to analysis of 10,500,000 samples, sustainability mutations have remained constant from January 1, 2020, to April 28, 2022, in France, Spain, Greece, Netherlands, England, New Zealand, Senegal, Russia and the USA are shown in Figure 1.

**Figure 1.**
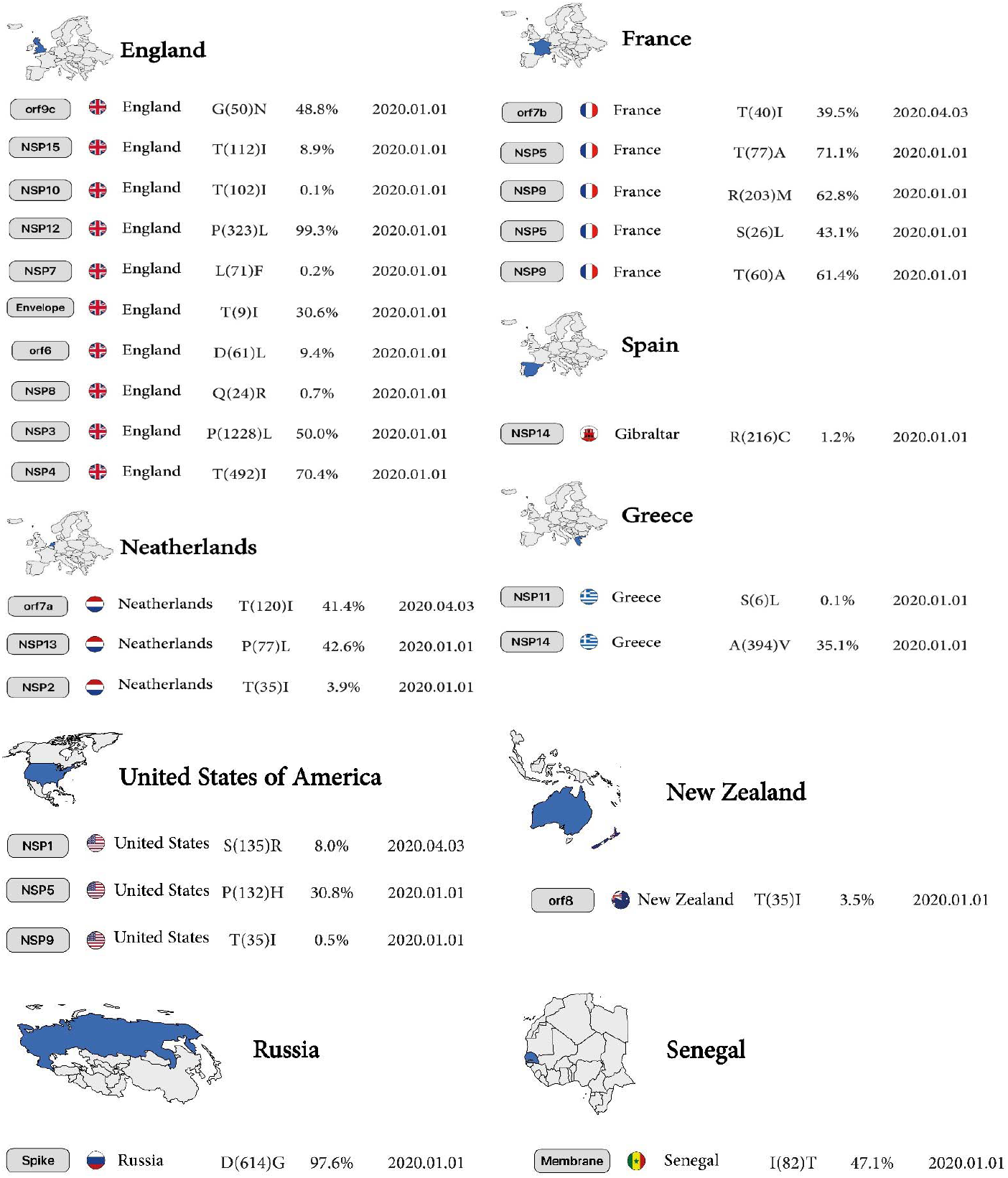
The first country reported sustainable mutation in each gene. From left to right, gene name, country name, Amino acid substitutions and positions, the Mutation frequency of all mutations occurring in that gene, and date of the first sample include mentioned mutation.

### NSP1

The highest mutation is the conversion of Serine to Arginine, the most common mutation in North America, with 8% reported in 2020. Subsequent mutations in the conversion of Glutamic acid to Aspartic acid and Histidine to Tyrosine are 0.8% and 0.2% in England and the United States, respectively. Finally, the tenth mutation that has remained stable over time is the conversion of Proline to Serine at 0.08%, reported on January 1, 2020, in England.

### NSP2

The conversion of Threonine to Isoleucine at 3.9% is the highest mutation in the Netherlands. The conversion of Valine to Isoleucine at 0.3% is the lowest mutation in North America in January 2020 for the nsp2 protein.

### NSP3

Mutations in the nsp3 protein include conversion of Proline to Leucine at a rate of 50% (highest mutation), and subsequent mutations including conversion of Proline to Serine, Alanine to Serine, Threonine to Isoleucine (position 183), Alanine to Aspartic acid, and Isoleucine to Threonine (position 1412) from France and England reported on January 1, 2020, and the conversion of Proline to Leucine (tenth mutation) is 5.1% in England reported on January 1, 2020.

### NSP4

The highest mutations in Europe were observed in England and the Netherlands, which include the mutations of Threonine to Isoleucine at 70.4% and the conversion of Valine to Leucine at 37.9%, respectively. The lowest mutation was detected in Northern Ireland with the replacement of Alanine to Valine at 0.5% for nsp4 protein reported on April 21, 2020.

### NSP5, NSP6

Conversion of Leucine to Phenylalanine (position 89) was observed in nsp5 protein with 1.4% reported on January 1, 2020, in North America; however, in New Zealand, a 2.9% AA change (position 37, mutant 5) was observed in the same month for nsp6 protein. In the nsp5 protein, conversion of Alanine to Valine (position 260, mutant 6) was observed at 0.2% in North America, but this mutation occurred (position 2, mutant 7) in South America at 1.1% for nsp6 protein in January 2020.

### NSP7

In the nsp7 protein, conversion of Leucine to Phenylalanine was observed in 2 different countries and positions, including England (position 71) at 0.2%, reported on January 1, 2020, and India (position 56) reported at 0.1%, reported on March 1, 2020 (Table1, Figure 2A).

**Table 1.**
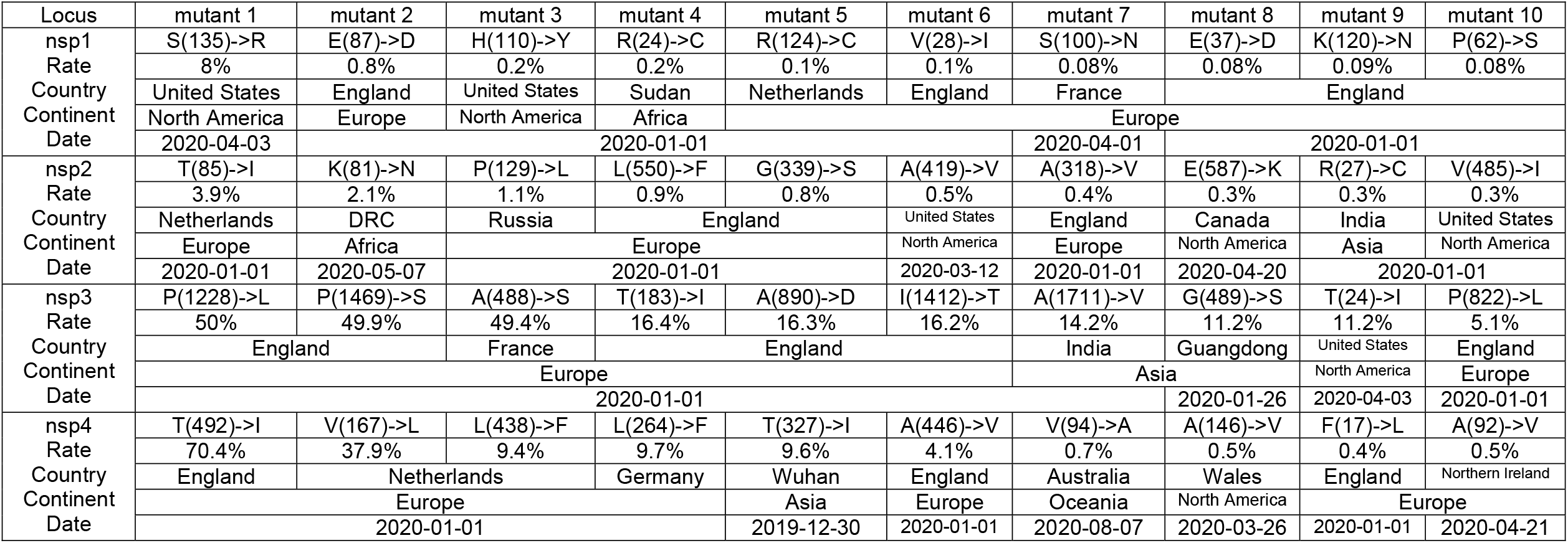
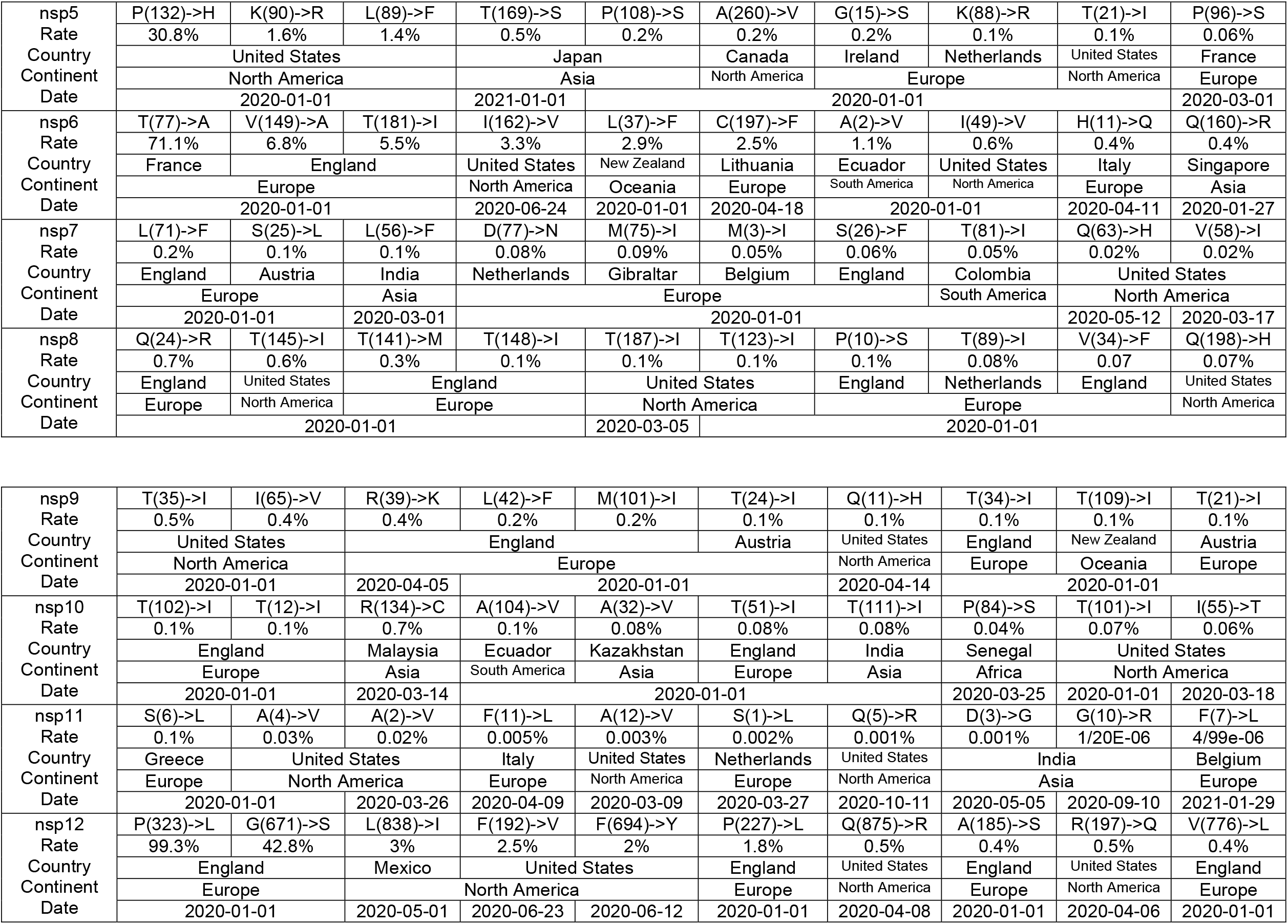
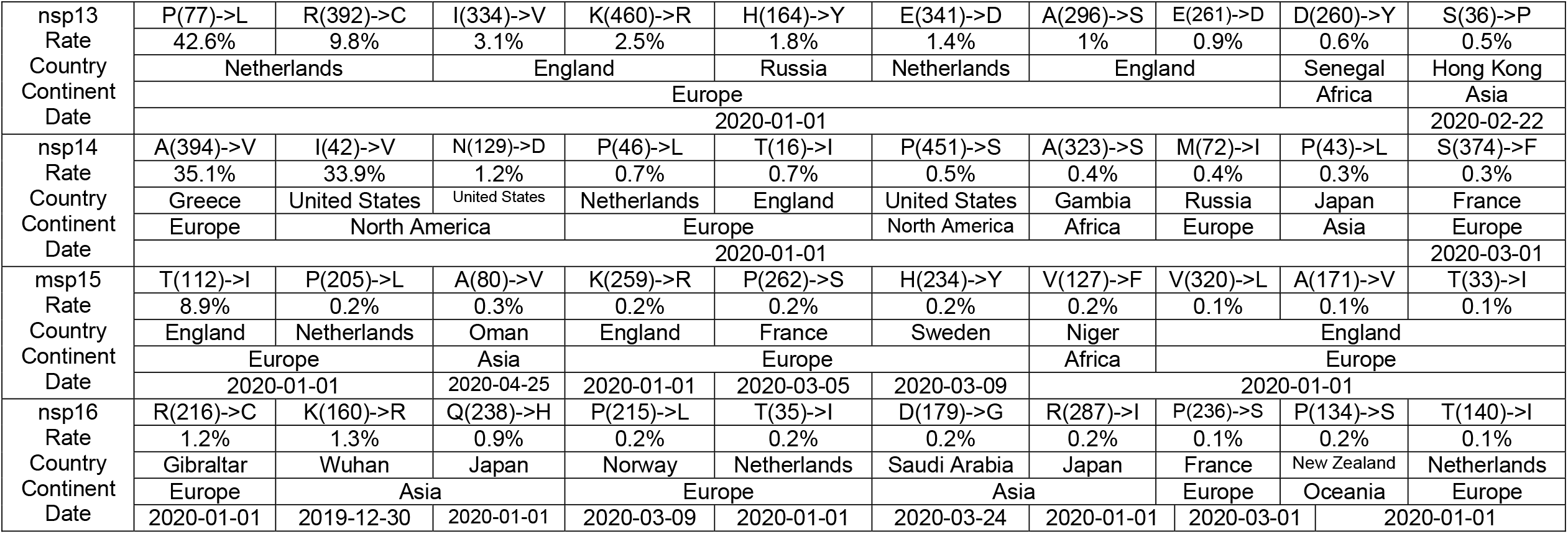
Top 10 of the First stable mutation in sars-cov2, including nsp1 to nsp16

**Figure 2A:**
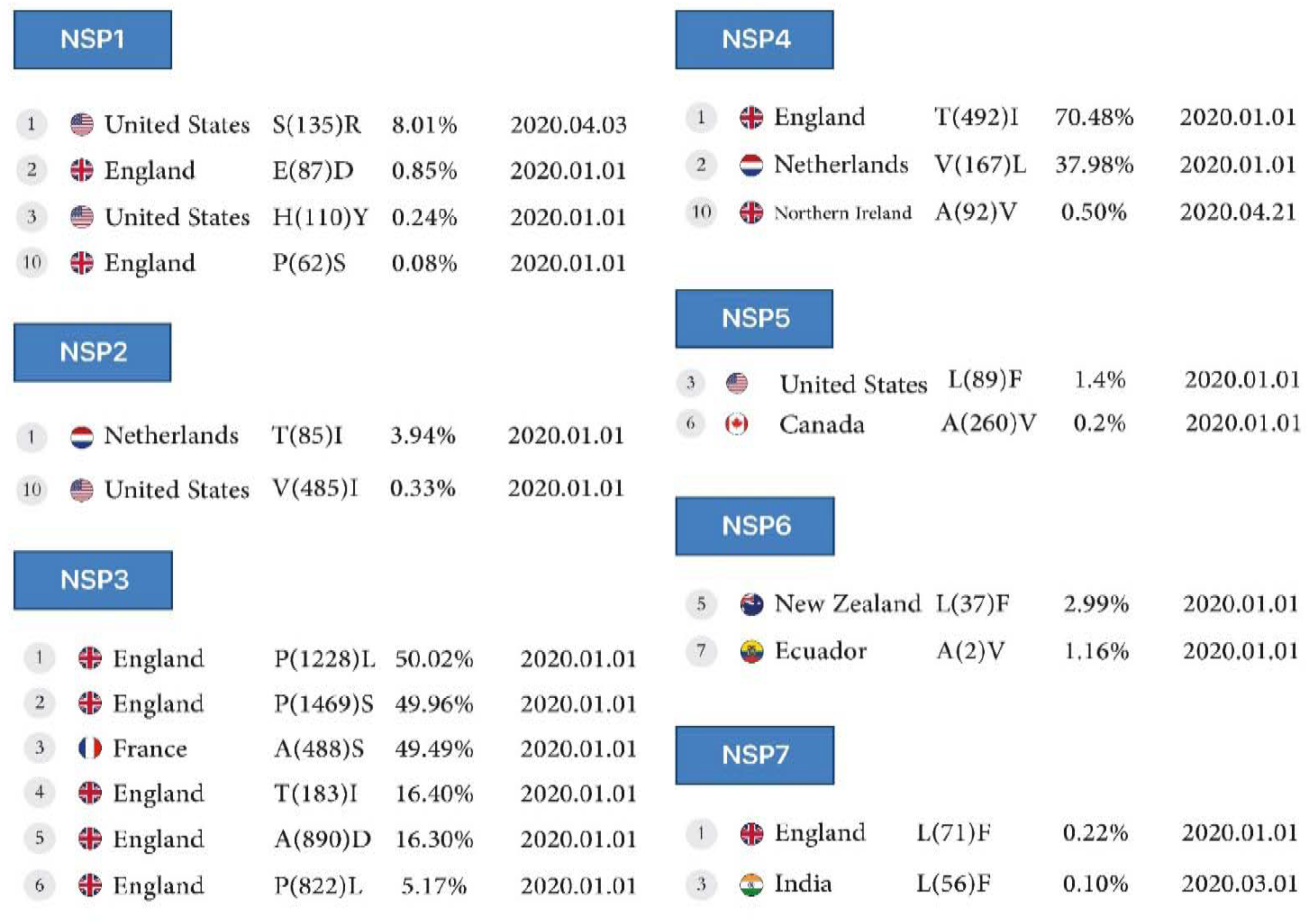
First stable mutations in SARS-CoV-2, including nsp1-nsp7

**Figure 2B:**
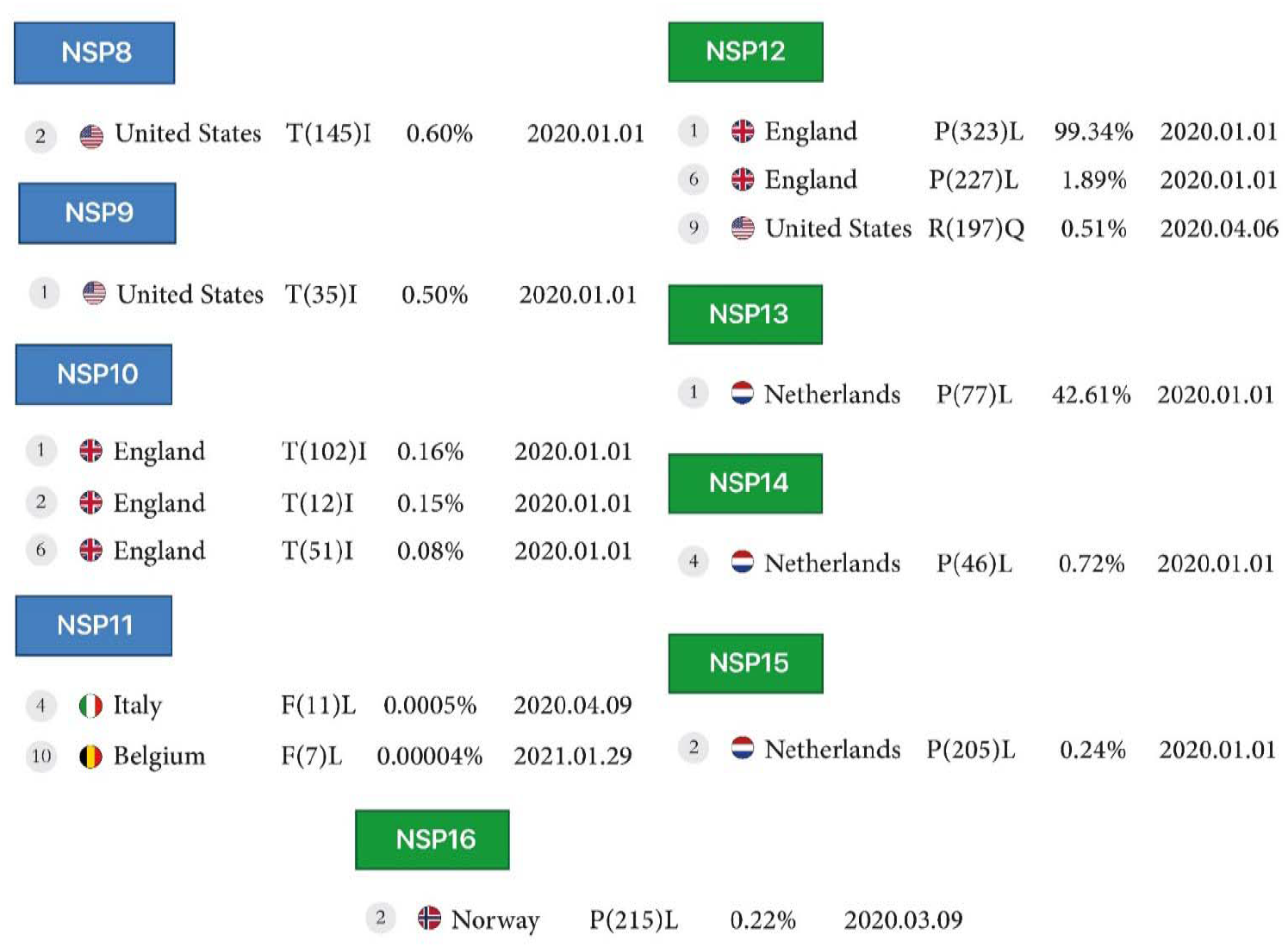
First stable mutations in SARS-CoV-2, including nsp8-nsp16

### NSP8, NSP9, NSP10

Conversion of Threonine to Isoleucine in North America for the nsp8 protein at 0.6% (position 145), for the nsp9 protein at 0.5% (position 35), and for the nsp10 protein at 0.1%, 0.1% and 0.08% (position 102, 12 and 51), respectively.

### NSP11

In the nsp11 protein, conversion of Phenylalanine to Leucine was observed in 2 different positions, 11 and 7, in Italy, reported on April 9, 2020, and in Belgium, reported on January 29, 2021.

### NSP12

The nsp12 protein mutations include the conversion of Proline to Leucine in 2 different positions, 323 and 227 at 99.3% and 1.8%, respectively, in England, and conversion of Arginine to Glutamine in position 197 at 0.5% and Glutamine to Arginine in position 875 at 0.5% in North America, all reported on April 6 and April 8, 2020.

### NSP13, NSP14, NSP15, NSP16

The conversion of Proline to Leucine in position 77 (the first mutation) at 42.6% for the nsp13 protein, and position 46 (the fourth mutation), at 0.7% for the nsp14 protein, and in position 205 at 0.2% for the nsp15 protein reported from the Netherlands on January 1, 2020, and in position 215 (the fourth mutation) at 0.2% for the nsp16 protein from Norway, reported on March 9, 2020 (Table1, Figure2B).

### ORF3a

In the Orf3a protein, the highest detected mutation is the conversion of Serine to Leucine at the rate of 43.1% reported in France, and the lowest mutation is the conversion of Tryptophan to Cysteine at the rate of 0.7%reported from Mexico, with both mutations occurring in early 2020. In North America, mutations were detected in the conversion of Threonine to Isoleucine at 10.2%, Glutamine to Histidine at 5%, and Glutamic acid to Glutamine at 2.7%, as reported in the USA. In South America, the conversion of Serine to Proline, at the rate of 1.2% in Brazil, was reported on March 9, 2020.

### ORF6

Four mutations were detected in England in early 2020, the conversion of Aspartic acid to Leucine, Proline to Leucine at 0.1%, and Valine to Phenylalanine at 9.4%, 0.1% for two first and 0.03% for the last. Also, another There was mutation of the conversion of Arginine to Serine at 0.05% was reported in the Netherland.

### ORF7a

The highest mutation of Orf7a was from the conversion of Threonine to Isoleucine at 41.4%, and Valine to Alanine at 41.1% found in the Netherlands and France, respectively, in early 2020. There was another mutation in England, the conversion of Threonine to Isoleucine at the rate of 0.2% found in England, the conversion of Leucine to Phenylalanine at the rate of 0.9% in France, and the conversion of Proline to Serine at the rate of 0.1% found in Wuhan. In Asia, there were two mutations: the conversion from Proline to Leucine at a rate of 1.5% in Saudi Arabia and Arginine to Glycine at a rate of 0.2% in India. The United States also had two mutations, the conversion of Valine to Isoleucine at the rate of 0.6% and Histidine to Tyrosine at 0.2%.

### ORF7b

The highest rate of mutation of Orf7b is the conversion of Threonine to Isoleucine at a rate of 39.5% found in France. There were two more mutations in England, the conversion of Serine to Leucine and Methionine to Isoleucine at the rate of 0.1% and 0.05%, respectively. Mutations in Europe also included the conversion of Histidine to Tyrosine at the rate of 0.1% found in France and Alanine to Serine at 0.08% found in Greece. Three mutations were found in Asia, the conversion of Alanine to Valine at the rate of 0.04% in Guangdong, Serine to Leucine at 0.09% in Japan, and Cysteine to Phenylalanine at the rate of 0.05% in Thailand.

### ORF8

The highest mutation rate in Oceania was the conversion of Serine to Leucine at 3.5% found in New Zealand. In Europe, England had the most mutations with the conversion of Glutamic Acid to Lysine at the rate of 2.6%, Alanine to valine at the rate of0.5%, and Valine to Leucine at the rate of 0.4%. In Italy, there was a mutation with the conversion of Serine to Phenylalanine at the rate of 0.5%, and in Gibraltar, a conversion of Alanine to Serine at the rate of 0.3% (Table2, Figure 3A).

**Table 2.**
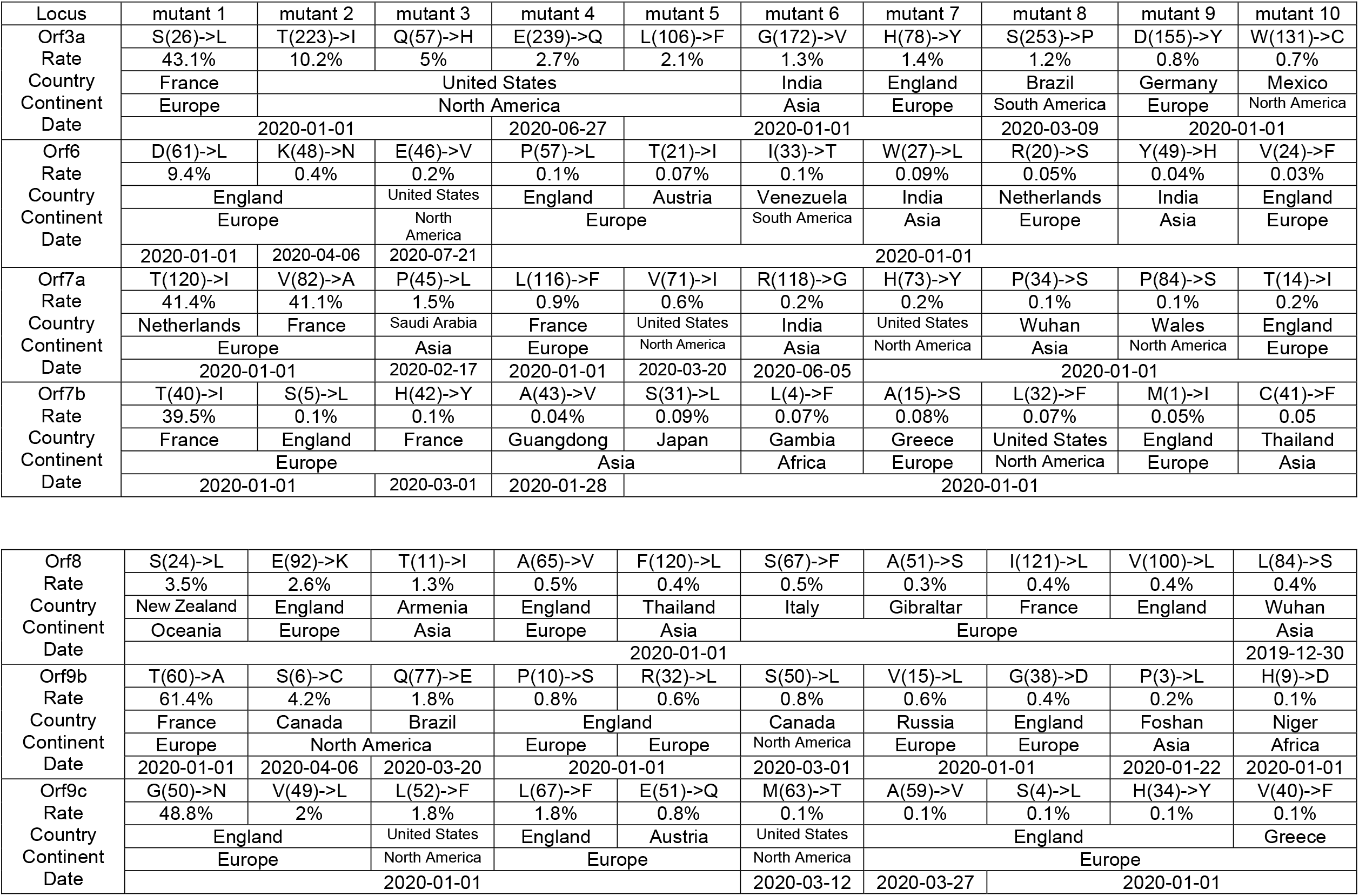
Top 10 of the First stable mutation in sars-cov2, including Orf3a to Orf9b

**Figure 3A:**
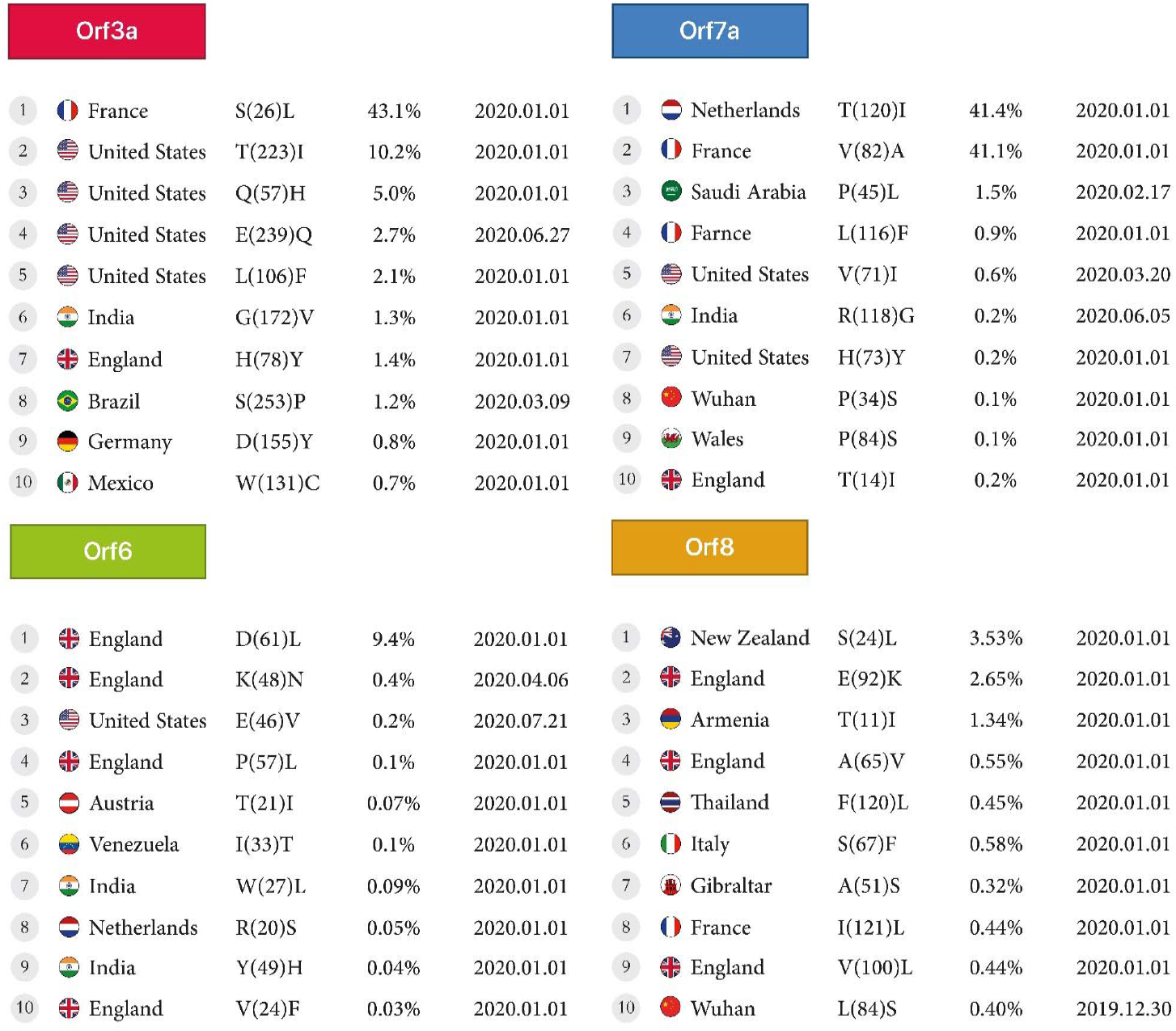
First stable mutations in SARS-CoV-2 including ORF3a, ORF6, ORF7a, ORF8

**Figure 3B:**
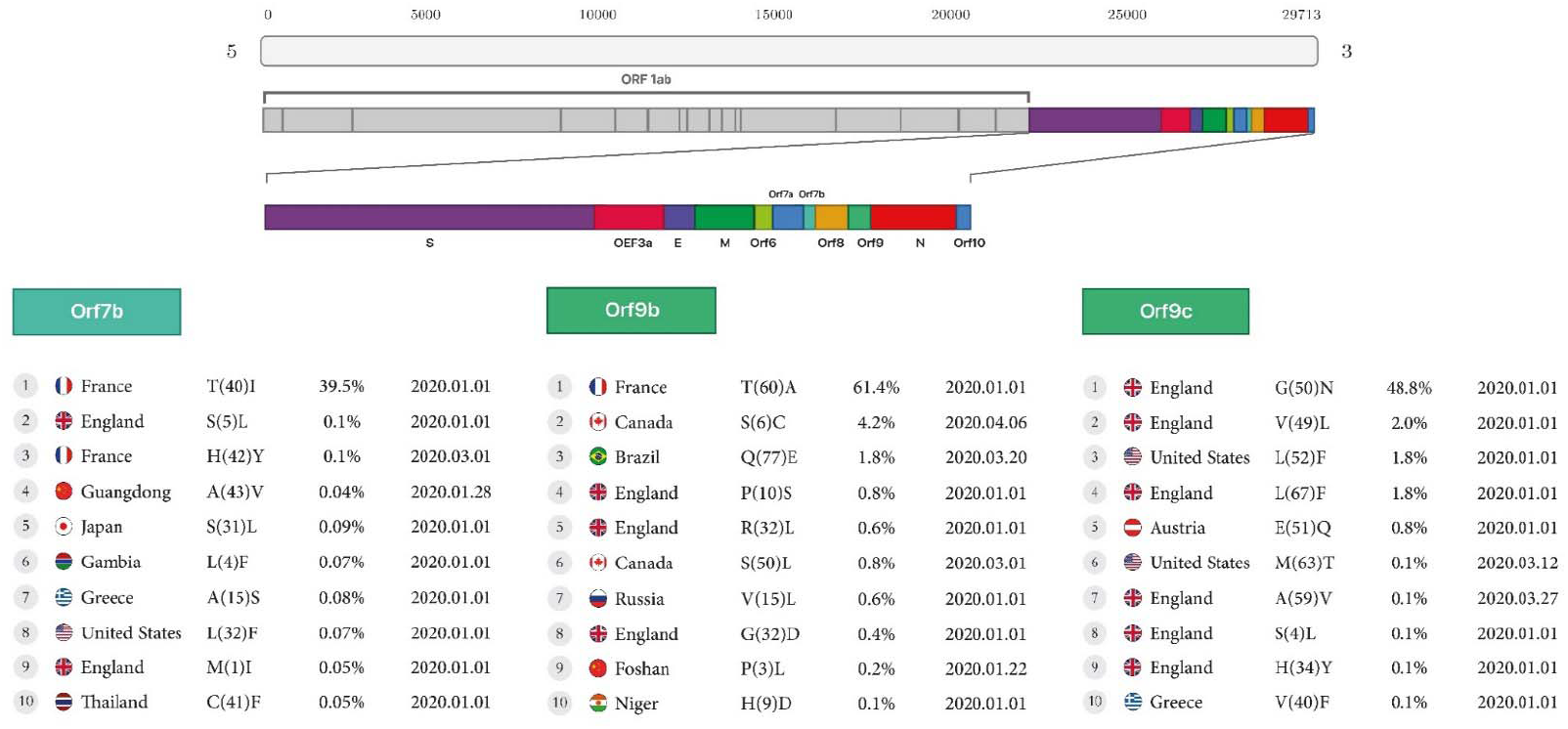
First stable mutations in SARS-CoV-2 including (ORF7b, ORF9b, ORF9c)

### ORF9b

The highest mutation rate in Orf9b is the conversion of Threonine to Alanine at the rate of 61.4% found in France, and additional mutations in Europe, were the conversion of Valine to Leucine at the rate of 0.6% in Russia, and Glycine to Aspartic acid at the rate of 0.4% in England. Two mutations were found in Canada, the conversion of Serine to Cysteine at the rate of 4.2% and Serine to Leucine at the rate of 0.8%. There was one mutation in Asia with the conversion of Proline to Leucine at the rate of 0.2%, found in Foshan.

### ORF9c

Europe had the most mutations in the Orf9c protein. There were seven mutations in England, the conversions of Glycine to Asparagine at the rate of 48.8%, Valine to Leucine at the rate of 2%, Leucine to Phenylalanine at the rate of 1.8%, Glutamic Acid to Glutamine at the rate of 0.8%, Alanine to Valine at the rate of 0.1%, Serine to Leucine at the rate of 0.1% and Histidine to Tyrosine at the rate of 0.1%. (Table2, Figure3B).

### Envelope, Membrane, Spike, Nucleoprotein

The conversion of Leucine to Phenylalanine was observed in 2 positions, 21 and 73, at 0.1% and 0.08% for the envelope protein, and the conversion of Serine to Phenylalanine was observed in mutant7 and mutant8 at 0.08% and 0.07% in England, reported on January 1, 2020. For the nucleoprotein, the conversion of Arginine to Methionine at position 203 at 62.8% was the highest mutation in France. The conversion of Alanine to Valine at position 220 at a rate of 2.5% was the lowest mutation found in England. Both were reported on January 1, 2020. For the membrane, the conversion of Isoleucine to Threonine at position 82, at a rate of 47.1%, was the highest mutation found in Senegal. The conversion of Serine to Asparagine in position 197 at a rate of 0.2% was the lowest mutation found in Spain. Both were reported on January 1, 2020. Finally, for the Spike protein, the conversion of Aspartic acid to Glycine at a rate of 97.6% was the highest mutation in Russia. The conversion of Aspartic acid to Tyrosine at a rate of 10% was the lowest mutation found in the Netherlands. Both were reported on January 1, 2020. (Table 3, Figure4).

**Table 3.**
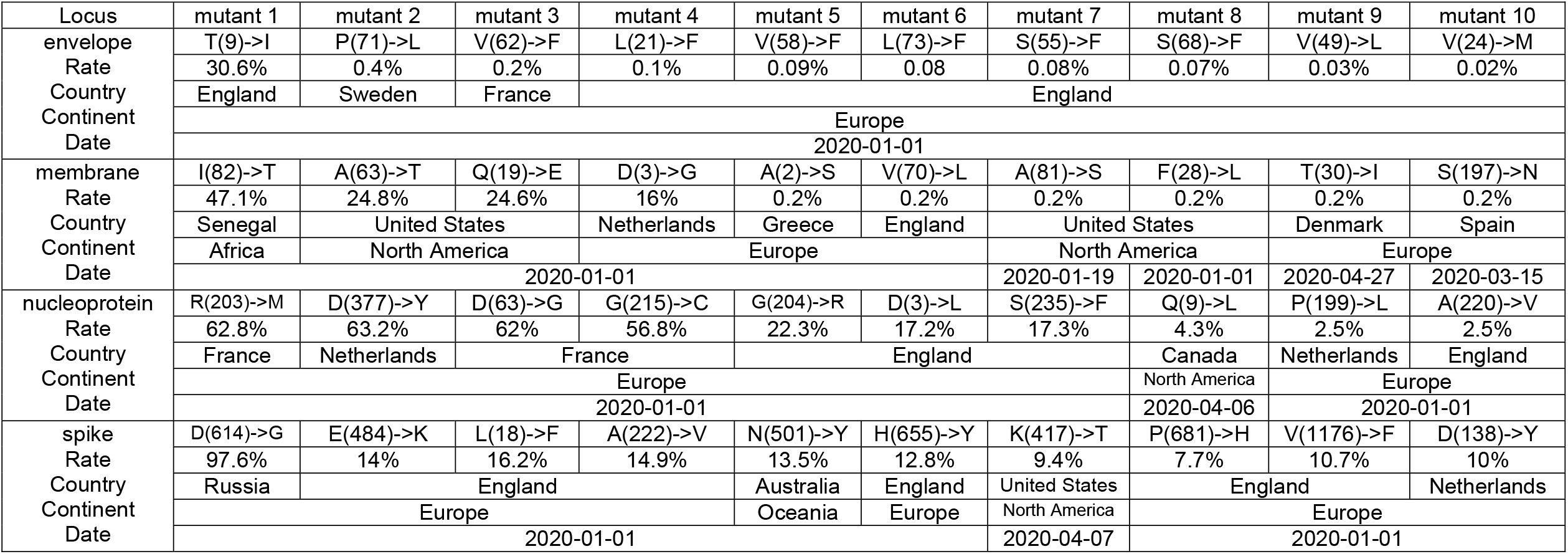
envelope, membrane, nucleoprotein, spike

**Figure 4.**
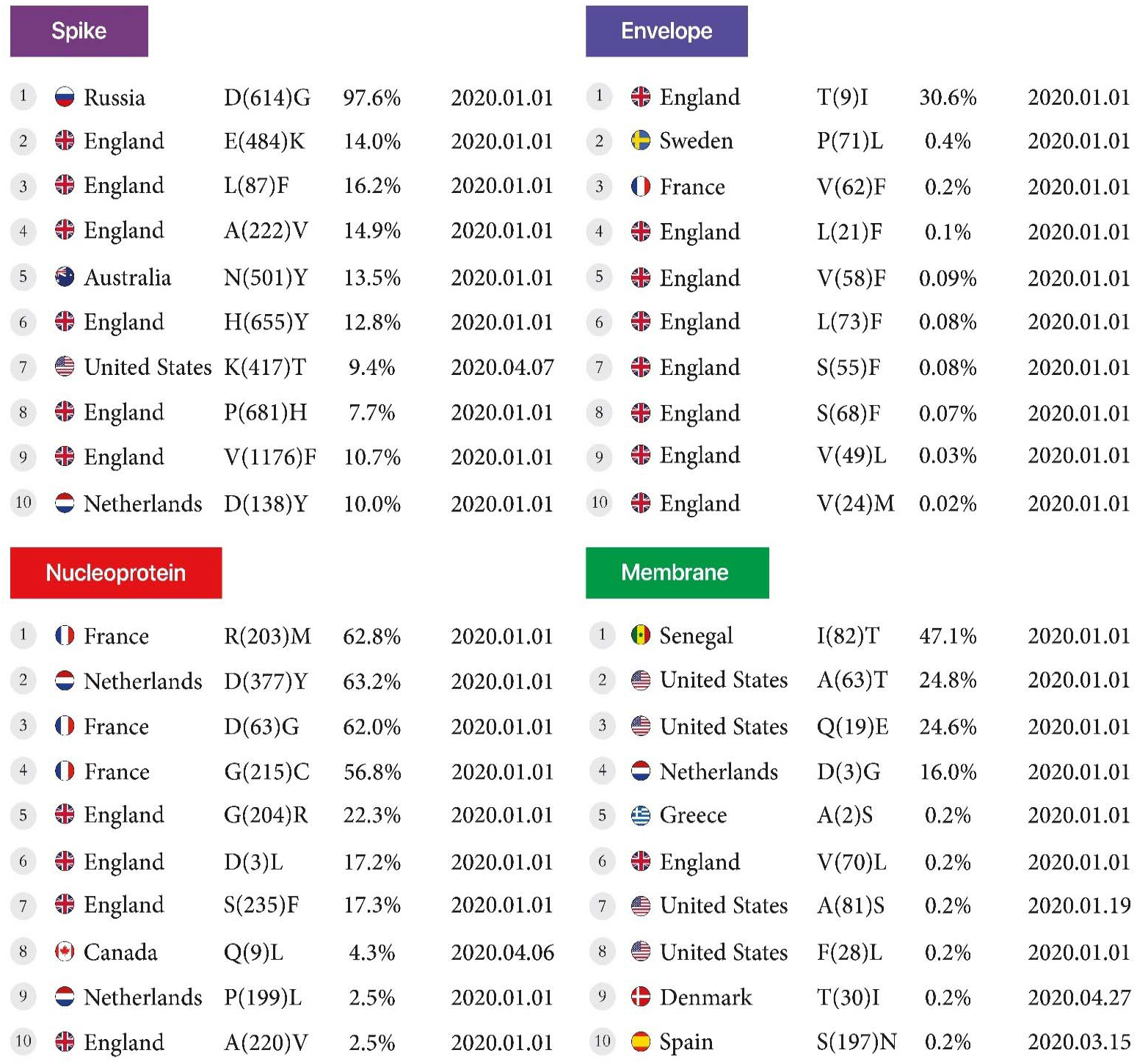
First stable mutation in sars-cov2 including Envelope, Membrane, Nucleoprotein, and Spike.

## DISCUSSION

During the early stages of the pandemic, the virus swiftly moved throughout the world and continued to spread globally. There was a more diversified mix of strains among the samples submitted to GISAID(14-16). Type G strains became more prominent in nations with higher resistance, resulting in a strain shift. Based on previous reports, the G strains may be a simpler target for vaccination because they contain larger quantities of surface proteins that target antibodies that bind to vaccines. Spike protein enables virus attachment to the human cell surface ACE2 receptor, allowing for viral entrance during infection(17-19). It is divided into two components called S1 and S2. The S1 unit has a receptor-binding domain (RBD) that may directly connect to the ACE2 receptor and the primary target of SARS-CoV-2 neutralizing antibodies (Ab). As a result, S1 is a hotspot for mutations with significant clinical significance in virulence, transmissibility, and host immune evasion(5, 20-22). The Alpha version features an N501Y mutation, which means that N asparagine has been replaced with Y tyrosine at the 501 residues, and K417N—lysine K has been replaced with asparagine N(23). A developing variety originating from B.1.1.7 also has the E484K mutation, which replaces glutamic acid E with lysine K. Other than N501Y, Beta and Gamma versions feature additional substitutions(23). The E484K mutation is present in the Beta variation, whereas the E484K and K417T mutations are present in the Gamma variant [GISAID-hCov19 Variants [Internet]. [cited 2021 June 6]. https://www.gisaid.org/hcov19-variants/]. The AA change D614G has been documented as a result of a nucleotide A to G mutation at position 23403 in the Wuhan reference strain. In early March 2020, this mutation was the first known alteration in the Spike analysis. During that period, the G614 form was uncommon but essential in Europe developments. The D614G mutation is frequently seen in conjunction with two additional mutations; C to T mutation at position 241 in the ‘5 UTR compared to the Wuhan reference sequence (silent C to T mutation at viral RdRp site) (P323L RdRp)(24). The V483A and G476S mutations were discovered in US samples initially, whereas the V367F mutant was found in China, Hong Kong, France and the Netherlands. The mutations V367F and D364Y have been shown to improve the stability of the spike protein structure and promote effective binding to the ACE2 receptor(11). A passenger returning to Taiwan from Wuhan reported an identical nucleotide-382 deletion in Orf8. However, the clinical consequence of this deletion has not been recorded. Orf8 is a good location for viral coronavirus alterations, according to research by Young E Barnaby et al. Deletion in this area appears to result in a milder illness with less systemic proinflammatory cytokine production and a more effective immune response to SARS-Cov2(25). Further research into this mutation might help to further therapy and vaccine development(25). NSP1 of Orf1a/Orf1ab and Orf8 have been identified as mutation hotspots linked to virulence and transmissibility. NSP1 is a critical protein that inhibits type I interferon activation in the host while promoting viral replication(23, 26). Yuan-Qin Min et al., 2021, concluded that in the nsp1 protein, the conversion of Aspartic Acid to Glutamic Acid occurred at position 75 in China, but in our study, the conversion of Glutamic Acid to Aspartic Acid occurred at position 87 in England. The present research is not consistent with our analyses(27). Orf8 is an immune-evasive protein that reduces the expression of major histocompatibility complex class I (MHC-I) in host cells(25). Recently, it was discovered that the Alpha variation found from a single immunocompromised person had a premature stop codon at position 27 of Orf8(10). S-protein mutation D614G has influenced the SARS-CoV-2 transmissibility rate due to increased affinity for olfactory epithelium. It has been shown to have higher transmissibility in animal models(28, 29). The Alpha variation has been reported to increase hospitalizations and death, which may be attributable to their resistance to neutralizing antibodies owing to RBD mutations(30). Yuan and colleagues discovered that the CR3022 neutralizing antibody derived from a SARS patient binds to the RBD region of the spike in the SARS-Cov-2 virus by producing the crystal structure of this complex. This crystal structure enables researchers to comprehend how the SARS-CoV-2 virus is discovered. Identifying and obtaining these epitopes enables designing a vaccine against the SARS-CoV-2 virus and the detection of antibodies to additional coronaviruses in the future(31). Wang and colleagues have identified 13 mutation sites in the SARS-CoV-2 Orf1ab, Orf3a, Orf8 and N regions, with 28144 in Orf8 and 8782 Orf1a showing a mutation rate of 30.53% and 29.47%, respectively(32). The SARS-CoV-2 RdRp (also known as nsp12) is a crucial component of the replication/transcription machinery. Compared to SARS-CoV, SARS-CoV-2 has a high homology for nsp12, suggesting that its function and mode of action may be well preserved(33). This is supported by a recent cryo-EM structural investigation of SARS-CoV-2 nsp12(34). RdRps are thought to be one of the most important targets for antiviral medication development against a wide range of viruses. Favipiravir, Galidesivir, Remdesivir, and Ribavirin are several RdRp inhibitors evaluated for SARS-CoV-2 treatment(35). Interestingly, the docking location is not near the RdRp catalytic domain(36). Furthermore, additional prospective medications like filibuvir, cepharanthine, simeprevir, and tegobuvir are expected to be RdRp inhibitors(37). Natural mutations in key residues for pharmacological effectiveness can result in drug resistance, with a considerable reduction in the binding affinity of these compounds to the RdRp(38-40). We found 10 mutations for Orf7a protein which, according to a recent study, showed that ectodomain Orf7a binds to monocytes in the human peripheral blood, reducing its ability to deliver antigens, inducing dramatic expression of proinflammatory cytokines, and since the lungs are the site of SARS-CoV-2 proliferation causing the accumulation of monocytes in the lungs(41). Also, the Orf8 signaling pathway promotes proinflammatory by activation of IL-17, which contributes to cytokine storm in SARS-CoV-2 infection(42), includes apoptosis(43) and antagonizes the IFN signaling pathway(44). It has been shown that Orf9b could cause the inflammasome’s activation, and significant antibody responses have been found to Orf9b. On the other hand, Orf9c expression impaired interferon signaling, antigen processing and presentation, complement signaling, and induced IL-6 signaling(45). Our study reports that the T60A mutation, which is correlated to Orf9b, is the more frequent mutation found in France, although Gopika et al. found this mutation in Thailand as well(46). Gupta S et al., concluded that in the Orf3a protein, Serine to Leucine conversion occurred at position 171 in India. However, we observed this AA conversion occurrence in France at a rate of 43.1%(47). Also, Michael D Sacco et al. concluded that mutations in Proline to Histidine at position 132, conversion of Leucine to Phenylalanine at position 89, and Lysine to Arginine at position 88 for the nsp5 protein in the omicron strain was consistent with our studies(48). In addition, Abdullah Shah et al. demonstrated that in the nsp2 protein, the AA change of Arginine to Cysteine occurred at position 197. The conversion of Valine to Isoleucine at position 378 occurred in Pakistan. However, in our study, the R27C mutation and the V485I mutation were detected in India and United States(49).

## Conclusion

The development of efficient therapy and early identification of variants rely on a better understanding of virus behavior, and forecast mutation jumps to control the worldwide pandemic. A geographical view and a survey on mutations are the keys to developing a strategic plan to prevent burst events likely to increase transmissibility, morbidity and mortality from the COVID19 pandemic. Recent findings of SARS-CoV-2 mutations call for a massive effort in the scientific community to identify new variants that may increase viral spreading and allow escape from natural or vaccine-induced neutralizing immunity. This study of SARS-CoV-2 variations may improve therapies, vaccine design, and diagnostic methods.

## MATERIALS AND METHODS

### Source of the sequences

The research analyzed data from the genome of SARS-CoV-2 amino-acid sequences (AASs). The AAS of the Wuhan-2019 virus, accession number ‘EPI ISL 402124’, was used as the reference sequence. This genetic sequence was applied to explore tightly correlated viruses against the whole database with BLASTn with a default parameter of 100. However, we obtained more than 100 results. This sample was used to compare against all 10,500,000 AASs samples obtained from the GISAID(14-16) database between 2020/1/1 - 2022/04/25. This database was provided by consent through Erasmus Medical Center’s permission.

### Analysis and processing of sequences

Python 3.8.0 was utilized to pre-process FASTA data and analyze sequence alignments and mutations. Each change between the sample and the reference marked a mutation, and the placement and substitution of the AAs were documented. Non-human samples, as well as samples containing unspecified AAs (reported as X), were excluded. The complete process was optimized using the ‘Numpy’ and ‘Pandas’ libraries.

### Detection of mutations

The algorithm design was based on all sequences having the same length. The terms ‘Refseq’ and ‘seq’ refers to reference sequence and sample sequence. The algorithm specification was:

for refitem,

seqitem in zip (refseq, seq)

If (refitem! = seqitem)

### Report of novel mutants

Following data extraction, the continent’s name and geographic locations of each sample were retrieved and presented using the pycountry-convert 0.5.8 program using Python ‘Titlecase’ package to create worldwide maps of mutation prevalence.

### Find candidate mutation

A high-frequency rate of at least 0.03% mutations was selected for plotting samples based on timelines.

### Statistical analysis

R 4.0.3 and Microsoft Power BI used data standardization and comparison chart outlining. Each region’s normalized frequency was presented to properly compare the data from each continent. For this purpose, the number of mutations on each continent was divided by the number of similar sequences in equal proportions.

## List Of Abbreviations

(SARS-CoV-2): Severe acute respiratory syndrome coronavirus 2
(ORF): Open reading frames
(NSP1): Nonstructural protein 1
(COVID-19): Coronavirus disease 2019
(AAS): Amino acid sequence
(AA): Amino acid
(IFN): Interferon

## Supplementary

- Supplementary file1 (first mutant).xlsx

## Declaration of competing interest

The authors declare that they have no conflicts of interest that might be relevant to the contents of this manuscript, and the research was carried out regardless of commercial or financial relationships that may cause any conflict of interest.

## Funding

This work was partially supported by the NIH grants and 2U54CA143727, 5P30GM114737, 5P20GM103466, 5U54MD007601, 5P30CA071789.

## Acknowledgments

The authors thank all of the researchers who have shared genome data openly via the Global Initiative on Sharing All Influenza Data (GISAID).

## Authors’ contributions

Conceptualization, M.M. and K.R, and Y.D; methodology, M.M and K.R; software, M.M and K.R; validation, M.M., K.R., S.T and D.L.K.; formal analysis, M.M, and K.R; investigation, M.M, S.T.H, A.G, M.M.S and A.F; resources, K.R, S.T, and A.M.N; data curation, M.M, K.R, A.M.N, S.T, and D.L.K; writing—original draft preparation, S.T.H, and A.G.; writing—review and editing, S.T, D.L.K.; visualization, A.M.N.; supervision, Y.D.; project administration, Y.D, M.M; funding acquisition, Y.D.

## Data availability

- The raw data supporting the conclusions of this article is available in supplementary file(s).

---

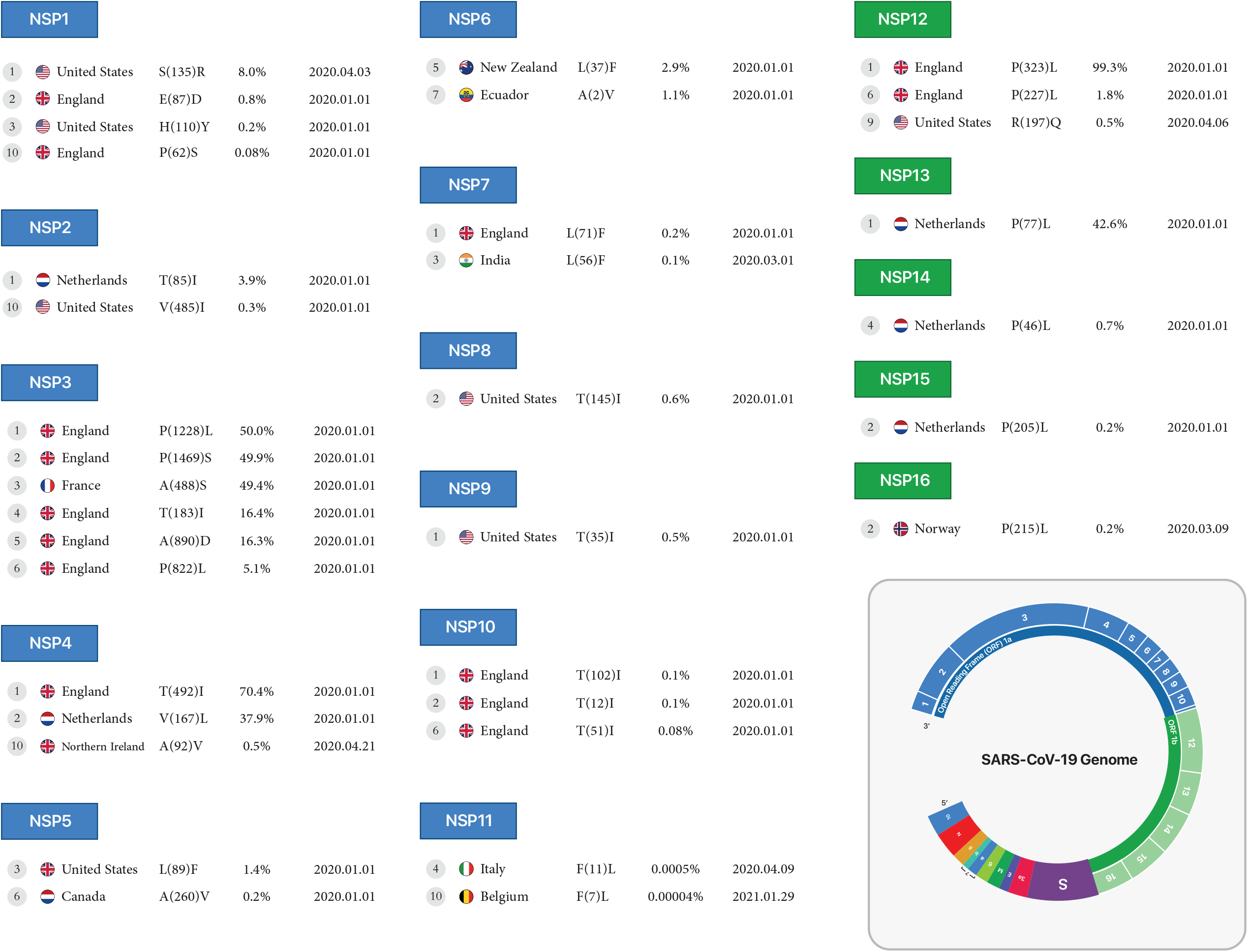

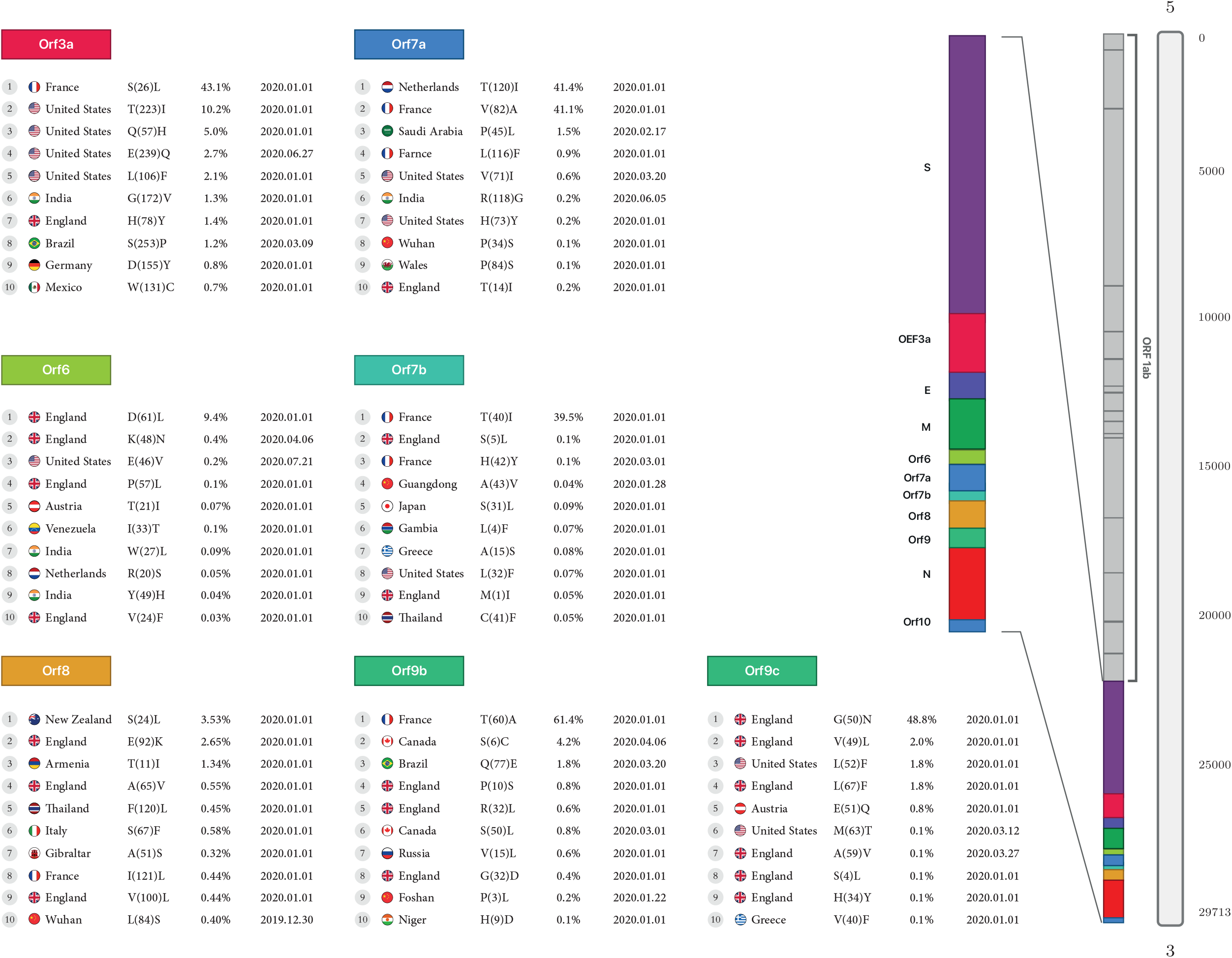

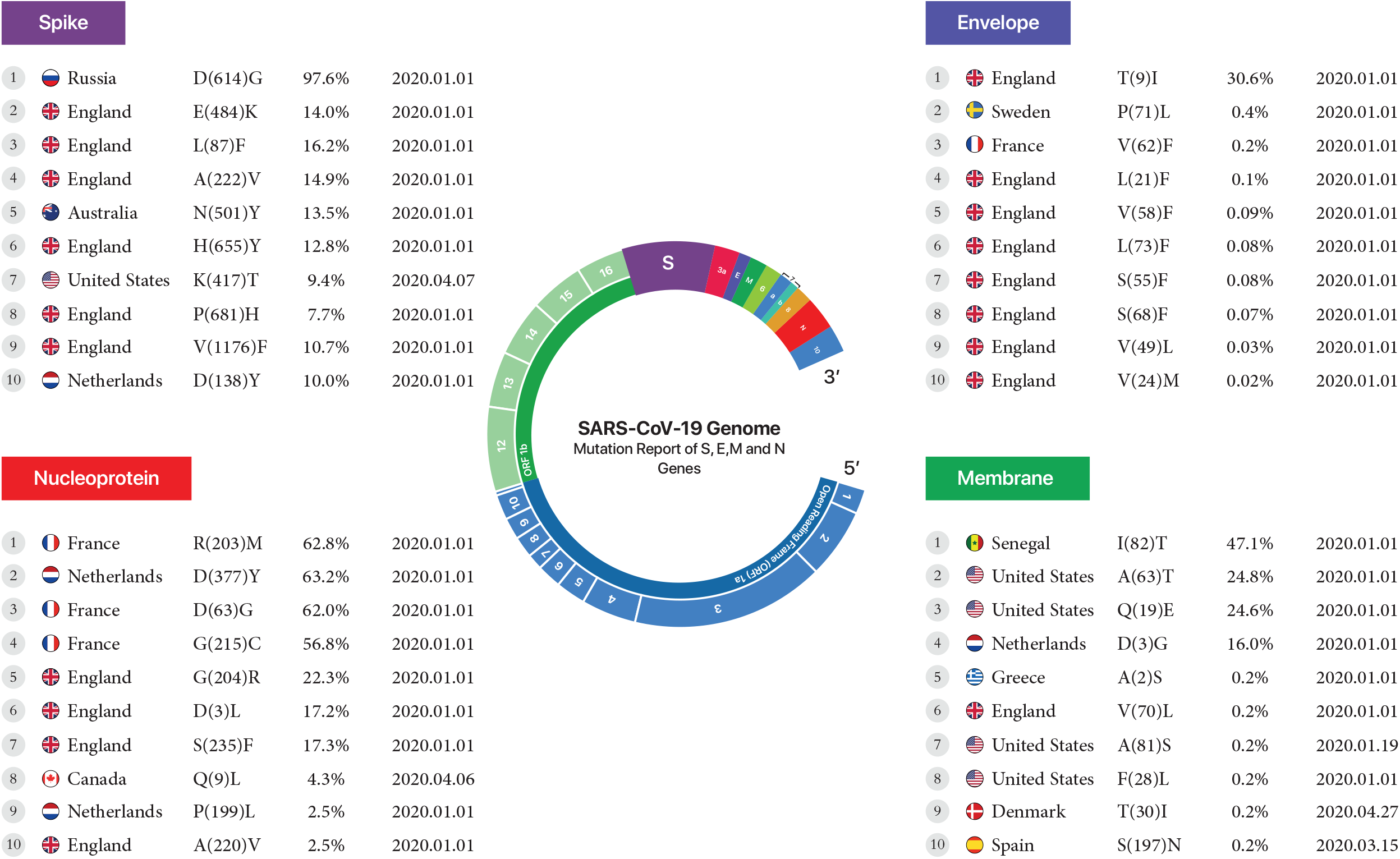

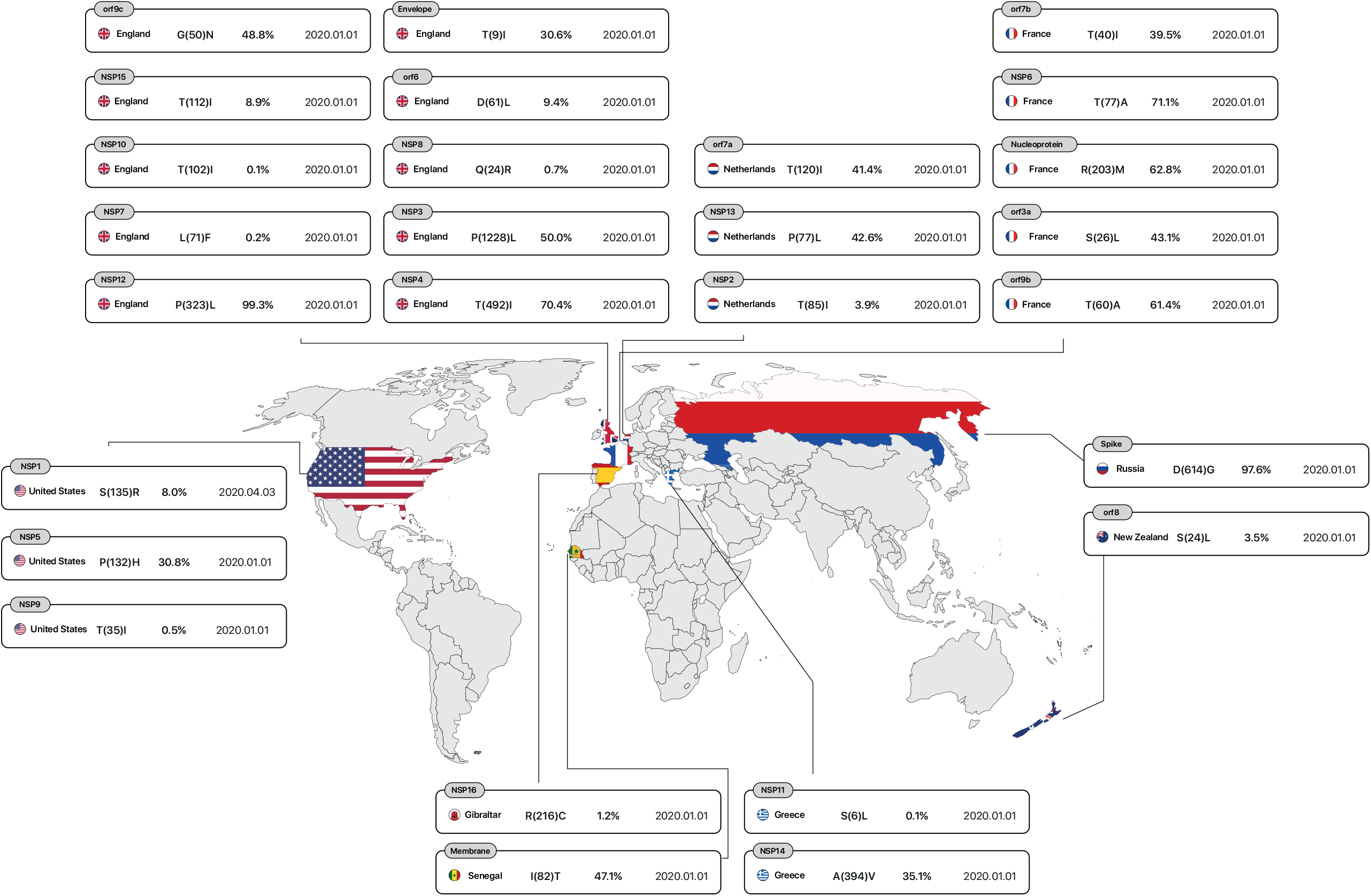

